# *Leptospira* enrichment culture followed by ONT Nanopore sequencing allows better detection of *Leptospira* presence and diversity in water and soil samples

**DOI:** 10.1101/2022.06.16.496521

**Authors:** Myranda Gorman, Ruijie Xu, Dhani Prakoso, Liliana C.M. Salvador, Sreekumari Rajeev

## Abstract

**Background:** Leptospirosis, a life-threatening disease in humans and animals, is one of the most widespread global zoonosis. Contaminated soil and water are the major transmission sources in humans and animals. Clusters of disease outbreaks are common during rainy seasons.

**Methodology/Principal Findings:** In this study, to detect the presence of *Leptospira*, we applied PCR, direct metagenomic sequencing, and enrichment culture followed by metagenomic sequencing on water and soil samples. Direct sequencing and enrichment cultures followed by PCR or sequencing effectively detected pathogenic and nonpathogenic *Leptospira* compared to direct PCR and 16S amplification-based metagenomic sequencing in soil or water samples. Among multiple culture media evaluated, Ellinghausen-McCullough-Johnson-Harris (EMJH) media containing antimicrobial agents was superior in recovering and detecting *Leptospira* from the environmental samples. Our results show that enrichment culture followed by PCR can be used to confirm the presence of pathogenic *Leptospira* in environmental samples. Metagenomic sequencing on enrichment cultures effectively detects the abundance and diversity of *Leptospira* spp from environmental samples.

**Conclusions/Significance:** The selection of methodology is critical when testing environmental samples for the presence of *Leptospira*. Selective enrichment culture improves *Leptospira* detection efficacy by PCR or metagenomic sequencing and can be used successfully to understand the presence and diversity of pathogenic *Leptospira* during environmental surveillance.

**Author Summary:** Leptospirosis, a life-threatening disease in humans and animals, is one of the most widespread global zoonosis. Contaminated soil and water are major sources of transmission in humans and animals. For this reason, clusters of disease outbreaks are common during the rainy season. In this study, *Leptospira* enrichment cultures followed by PCR and sequencing detected pathogenic and nonpathogenic *Leptospira* in soil and water samples. The pathogenic and intermediate groups of *Leptospira* were more prevalent in soil samples tested. Metagenomic sequencing on enrichment culture is effective in detecting the abundance and diversity of various *Leptospira spp*. in environmental samples. Soil samples in proximity to water may be an ideal niche for *Leptospira* growth and survival and may be an appropriate sample of choice for testing.

## Introduction

Many species of *Leptospira*, a spirochete bacterium that causes leptospirosis, are maintained in the renal tubules of numerous mammalian species and the environment (1). Leptospirosis is a life-threatening illness in humans, causing approximately 1 million cases and 60,000 deaths annually (2). A variety of mammals following *Leptospira* infection may become clinically ill or remain as asymptomatic renal reservoirs of infection. They shed bacteria through the urine and act as the source of infection to other animal hosts and environmental contamination (3). Leptospirosis is endemic to tropical countries, and outbreaks occur during natural disasters where humans come into contact with the contaminated environment. The environmental route is the most common mode of *Leptospira* transmission in humans. The host and the environment interface play a major role in the epidemiology and transmission of *Leptospira* infection. In addition to sporadic outbreaks during recreational water activities, large clusters of outbreaks after severe rain and flooding are more common in tropical countries. Continuous changes in climatic landscapes might increase the number of outbreaks occurring globally. A critical gap in knowledge on environmental persistence and cycling of *Leptospira* needs to be addressed (4). A number of studies have been conducted to investigate the level and type of *Leptospira* commonly found in environmental samples by applying multiple techniques.

The sensitivity and specificity of *Leptospira* detection in environmental samples can be complicated by the coexistence of chemical, physical and biological contaminants. Low levels of *Leptospira* present in the environmental sample among abundant contaminant microorganisms can also lead to false-negative results. Therefore, improvements in methods are needed for the accurate detection of *Leptospira* in environmental samples. Recently with the advent of Next Generation Sequencing (NGS) methods, the assessment of the microbial composition of environmental samples for disease surveillance has become a routine practice. For example, Oxford Nanopore Technologies (ONT) technology has been widely used for disease and environmental surveillance (5-8). We propose combining traditional selective culture methods with advanced sequencing could improve the *Leptospira* detection in the environmental samples. In this study, we evaluated multiple methods including selective enrichment culture, direct PCR, 16S rRNA gene amplification based sequencing, direct metagenomic sequencing, and *Leptospira* enrichment culture followed by metagenomic sequencing to detect the presence of *Leptospira* DNA in environmental samples.

## Materials and methods

### Sample Collection and processing

We collected representative soil and water samples from a local creek where abundant human and animal activity was observed. We collected one liter of water and approximately 50 g of soil from the damp edge of the creek from where water was collected in sterile containers and were transported to the laboratory on ice. After mixing the water thoroughly, we added 10 mL of Ellinghausen-McCullough-Johnson-Harris (EMJH) liquid medium (Becton Dickinson, Sparks, MD, USA) supplemented with Difco *Leptospira* Enrichment EMJH (Becton Dickinson, Sparks, MD, USA) media to the top of the water sample to enrich and attract *Leptospira*. After settling the sample for three hours, 200 mL of water from the top was collected and filtered with a 40 µm nylon filter. The filtrate was then divided into two 100 mL aliquots. We spiked one of the 100 mL aliquots with *Leptospira interrogans* serovar Copenhageni (10^7^ bacteria per mL) to use as the control, and the second aliquot was designated as the test sample. The samples were further divided into 50 mL aliquots for PCR, sequencing, and culturing.

For the processing of soil samples, the 25 g of soil was divided between two flasks and then mixed with 100 mL of phosphate-buffered saline (PBS). After mixing thoroughly for five minutes, the sample was allowed to settle for thirty minutes. Then 100 mL of EMJH media was added to the top of the samples and allowed to settle overnight. A longer settling time was required to obtain a cleaner sample for inoculation. Once settled, 80 mL of supernatant from each flask was collected and filtered through a 40 µm nylon filter. The filtrate was then aliquoted into two 75 mL samples. We spiked one of the aliquots with *Leptospira interrogans* serovar Copenhageni (10^7^ bacteria per-mL) and designated it as “control”. The non-spiked sample is designated as “test “sample. A schematic diagram showing soil and water processing is shown in supplemental figures 1 A and B.

**Figure 1:**
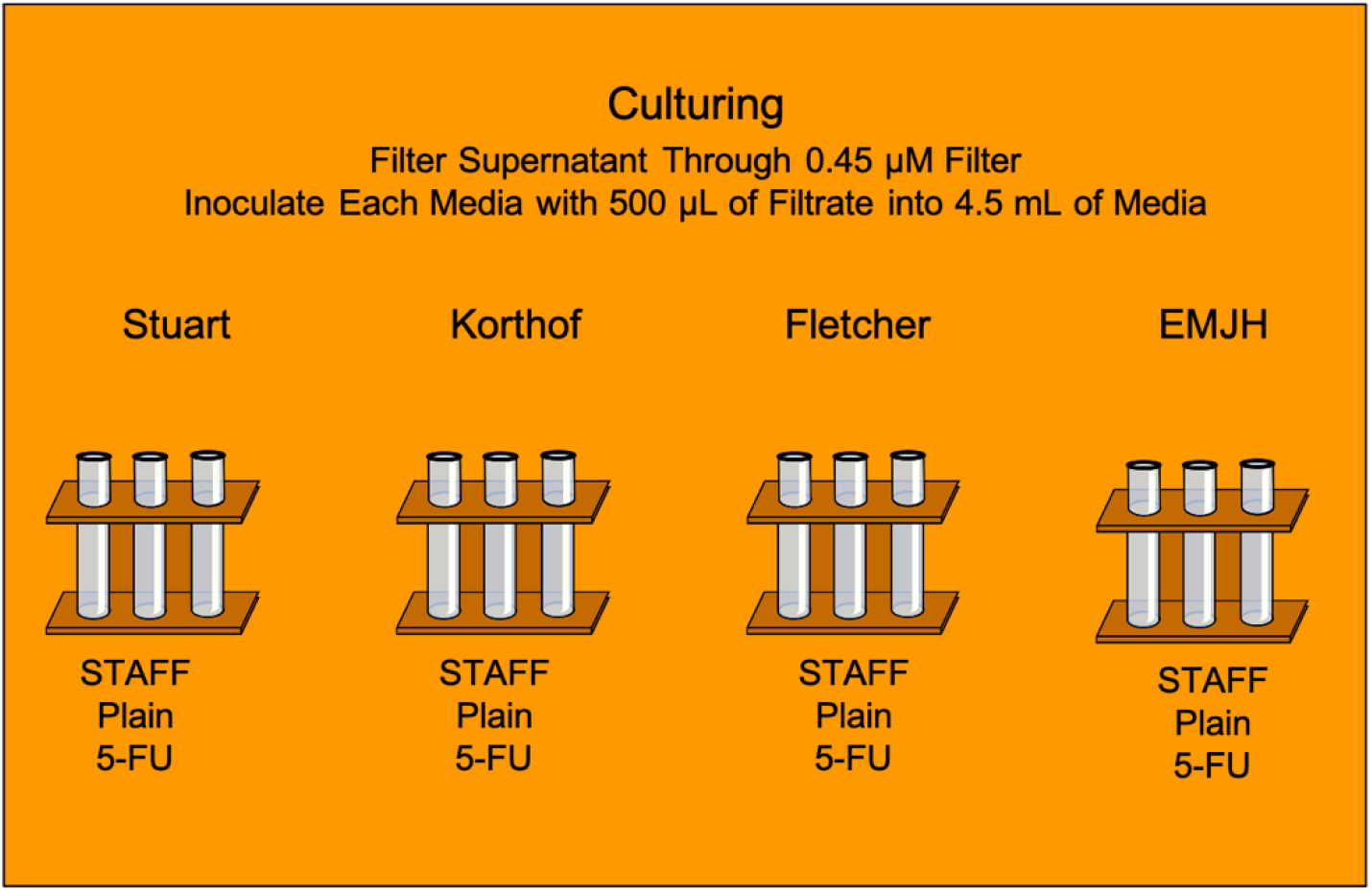
A schematic representation of sample inoculation for culture.

### *Leptospira* detection using direct PCR from water and soil

The 50 mL aliquots of test (non-spiked) and control (spiked) samples were centrifuged at 4,000 x g for forty minutes. The pellet was collected and then reconstituted with 10 mL PBS. Then 1 mL aliquots were pipetted into ten 1.5 mL collection tubes and stored at -20 ºC. DNA was extracted from three replicates of the spiked and test samples using the Quick-DNA Fecal/Soil Microbe Miniprep Kit (Zymo Research, Irvine, CA, USA) following the manufacturer’s protocol. The extracted DNA samples were then tested with Real-Time PCR targeting genes *LipL32, 16S rRNA, and 23S rRNA* to confirm the presence of *Leptospira* DNA (9-11) using a Q^®^ Quantabio (Quantabio, Beverly, MA, USA) thermocycler. The cutoff for a positive sample was set at a Cq value of 40.

### 16SrRNA gene-based metagenomic sequencing

This procedure was performed following a recent publication describing monitoring fresh water for pathogens (12)Briefly, extracted DNA samples were amplified using the full length of *16S rRNA* gene primers with common primer binding sequences 27f and 1492r, attached to unique 24 bp barcodes and nanopore motor protein tether sequence. The PCR was performed with 600 nM of each forward and reverse primer, 25 µL of Premix Taq DNA Polymerase (TakaraBio, Shiga, Japan), and a 10 µL DNA template in a 50 µL reaction. The amplification cycles used the following conditions 94 °C for 2 minutes, followed by 35 cycles of 94 °C for 30 seconds, 60 °C for 30 seconds, and 72 °C for 45 seconds, with final elongation at 72 °C for 5 minutes. The amplicons from the PCR step were purified using NucleoSpin Gel and PCR Clean-up (Macherey Nagel, Duren, Germany) following the manufacturer’s protocol. The barcoded amplicon samples were pooled in equimolar ratios, and library preparation and sequencing were conducted using Ligation Sequencing Kit SQK-LSK-109 (Oxford Nanopore Technologies, Oxford, UK) on the MinION (Oxford Nanopore Technologies, Oxford, UK) sequencing platform following the manufacturer’s instructions.

### Metagenomic sequencing directly from the environmental samples

The samples were spun down at 4,000 x g for forty minutes, and the supernatant was discarded, leaving a pellet in 10 mL of supernatant. After thorough mixing, 1 mL was aliquoted into ten 1.5 mL microcentrifuge tubes, then centrifuged at 14,000 x g for three minutes, and the supernatant was removed from each tube, leaving 200 µL with the pellet. DNA was extracted using Quick-DNA Fecal/Soil Microbe Miniprep Kit (Zymo Research, Irvine, CA, USA) according to the manufacturer’s instructions. The extracted DNA from water and soil underwent a further purification step using Monarch^®^PCR & DNA Cleanup Kit (New England Biolabs, Ipswich, MA, USA). DNA Library preparation was conducted using Native barcoding genomic DNA Kit SQK-LSK 109 combined with EXP-NBD104 (Oxford Nanopore Technologies, Oxford, UK) following the manufacturer’s instructions. After the end repair step, DNA from samples was barcoded and pooled in equimolar amounts to make one library, followed by adapter ligation and sequencing for approximately 48 hours on the MinION (Oxford Nanopore Technologies, Oxford, UK) sequencing platform.

### *Leptospira enrichment* culture, followed by PCR and metagenomic sequencing

We tested multiple media and antimicrobial combinations to enrich and grow *Leptospira* from environmental samples. We used four commonly used *Leptospira* culture media Stuart, Korthof, Fletcher, and Ellinghausen–McCullough–Johnson–Harris (EMJH) media. We tested each of these media with (plain) and without the addition of antimicrobials, 5-Fluorouracil (5-FU) and an antimicrobial cocktail (STAFF) to control the growth of competing bacteria in the cultures (13). We inoculated 500 µL of the processed water and soil to each of these media. A schematic representation of the inoculation of soil and water is shown in Figure 1.

The cultures were then incubated in a 29 °C incubator for four weeks, monitored at 24 hours, 72 hours, and then once a week for four weeks using dark field microscopy (DFM). The samples with the presence of organisms exhibiting *Leptospira*-like motility and morphology were presumptively identified as positive for *Leptospira* and scored from 0 to +4 rating system based on the number of spirochetes present (Table 1).

**Table 1:** The scoring system used in this study to evaluate cultures.

| Scoring | The relative number of *Leptospira*-like organisms under DFM |
| --- | --- |
| 0 | None seen |
| +1 | Less than 25 |
| +2 | Between 25 and 50 |
| +3 | Between 50 and 100 |
| +4 | More than 100/too numerous to count |

The presence and level of other contaminating bacteria were also recorded at each time point of evaluation. After four weeks of incubation and monitoring, 1 mL from each culture presumptively identified to contain *Leptospira*-like bacteria were collected, and DNA was extracted (Zymo Quick-DNA Miniprep kit, Zymo Research, Irvine, CA, USA). The DNA was then tested by PCR using *LipL32, 16S rDNA, and 23S rDNA* primers as described above. To evaluate the composition and *Leptospira* diversity of the culture samples, we pursued metagenomic sequencing using DNA extracted from culture samples. A composite of positive samples of culture and soil was used to reduce the cost of testing. Briefly, extracted DNA was purified using SparQ PureMag Beads (Quantabio, Beverly, MA, USA) following the instruction from the manufacturer. The Native barcoding genomic DNA Kit SQK-LSK 109 combined with EXP-NBD104 (Oxford Nanopore Technologies, Oxford, UK) was used for library preparation. The samples underwent end-repair, barcode ligation for multiplexing, and adapter ligation and sequencing. The DNA sequencing was conducted using the MinION (Oxford Nanopore Technologies, Oxford, UK) sequencing platform for approximately 24 hours.

#### Sequence Analysis

All scripts used for sequence data analysis is available at: https://github.com/rx32940/Environmental_Lepto_detection. All samples were base called using Guppy v. 6.1.1 with High Accuracy setting (https://community.nanoporetech.com). Samples were demultiplexed using Porechop v. 0.2.4. (https://github.com/rrwick/Porechop). To trim customized barcodes and adapters from each read during demultiplexing, customized barcodes and adapters were added to Porechop’s Adapter.py file before demultiplexing. The command ‘— discard_middle” was specified to remove chimeric reads attached by two different barcodes. The quality of the filtered reads was assessed using NanoStat v 1.5.0 (14) and visualized using Pistis v 0.3.4 (https://github.com/mbhall88/pistis).

### Microbial composition profiling and *Leptospira* classification from 16S dataset

Since the length of bacterial 16S rRNA is around 1.5 kbp ^1^, reads smaller than 1.4 kbp and larger than 1.6 kbp were filtered using NanoFilt v. 2.8.0 (14) to remove potential existing contaminations. To classify each read’s microbial taxon, each sample’s filtered reads were mapped against SILVA v. 138.1 16S rRNA database (15) using Minimap2 v 2.17 (16) with the recommended option for Nanopore reads “-ax map-ont”. Statistics for the percentage of reads mapped to the database were assessed using the “stat” function in Samtools v.1.10 (17). Mapped Bam files were converted to Bed format using “sam2bed” function in BEDOPS v 2.4.39 (18) for the downstream analysis. Microbial composition and abundance for each sample were analyzed using R. Reads mapped to more than one microbial taxa were assigned to the lowest common ancestor (LCA) of all mapped taxa. Reads that could not be assigned to at least a family-level taxon were removed from the downstream analysis due to low discrimination. Reads classified under all the taxa belong to the same bacterial family were summarized to obtain each sample’s microbial composition at the family level. The microbial composition for each sample was summarized and visualized using “dplyr” (https://dplyr.tidyverse.org, https://github.com/tidyverse/dplyr) and “ggplot2” packages in R (https://ggplot2.tidyverse.org)

All reads mapped under phylum *Spirochaete* were extracted from each sample’s sequences file using the “subseq” function in SEQTK v. 1.2 (https://github.com/lh3/seqtk) using read ID’s. Extracted *Spirochaetota* reads were aligned with all *Leptospira* 16S rRNA sequences deposited in NCBI using MUSCLE v 3.8.0(19) and built neighboring joining (NJ) phylogeny using the “-maketree” option in MUSCLE v 3.8.0 for genetic relatedness evaluation. NJ phylogenies were visualized using the “ggtree” package in R(20).

### Microbial composition profiling and *Leptospira* classification and identification from direct Sequencing and Sequencing from the enrichment culture

Each sample’s microbial composition was profiled using Kraken2 v. 2.0.9 (21) with the maxikraken2 database (https://lomanlab.github.io/mockcommunity/mc_databases.html) using the default settings. Profiling results of all samples were combined into a single file using KrakenTools v 1.2 (https://github.com/jenniferlu717/KrakenTools). Microbial reads classified under each taxon were analyzed and summarized in R using “dplyr” package (https://dplyr.tidyverse.org, https://github.com/tidyverse/dplyr) and visualized using “ggplot2” (https://ggplot2.tidyverse.org) package. All reads mapped under *Leptospira* taxa were subset from microbial profiles of each sample to visualize the relative percentage of *Leptospira* species identified from each sample.

## Results

### Direct PCR results from environmental samples

For the direct real-time PCR, both the test and spiked (control) water and soil samples were tested using the *Leptospira* specific *16S rRNA, Lipl32*, and *23S rRNA* gene markers. The *16S rRNA* gene was amplified from all the samples, however, Cq values were high in the test samples suggesting low levels of *16S rDNA* (Figure 2). *Lipl32* amplification product was detected only in the control samples and not in the test samples. The amplification pattern of the *23S rRNA* gene was inconsistent and was detected in the soil test and water control samples, but not in the soil control and water test samples. PCR results are shown in Figure 2

**Figure 2:**
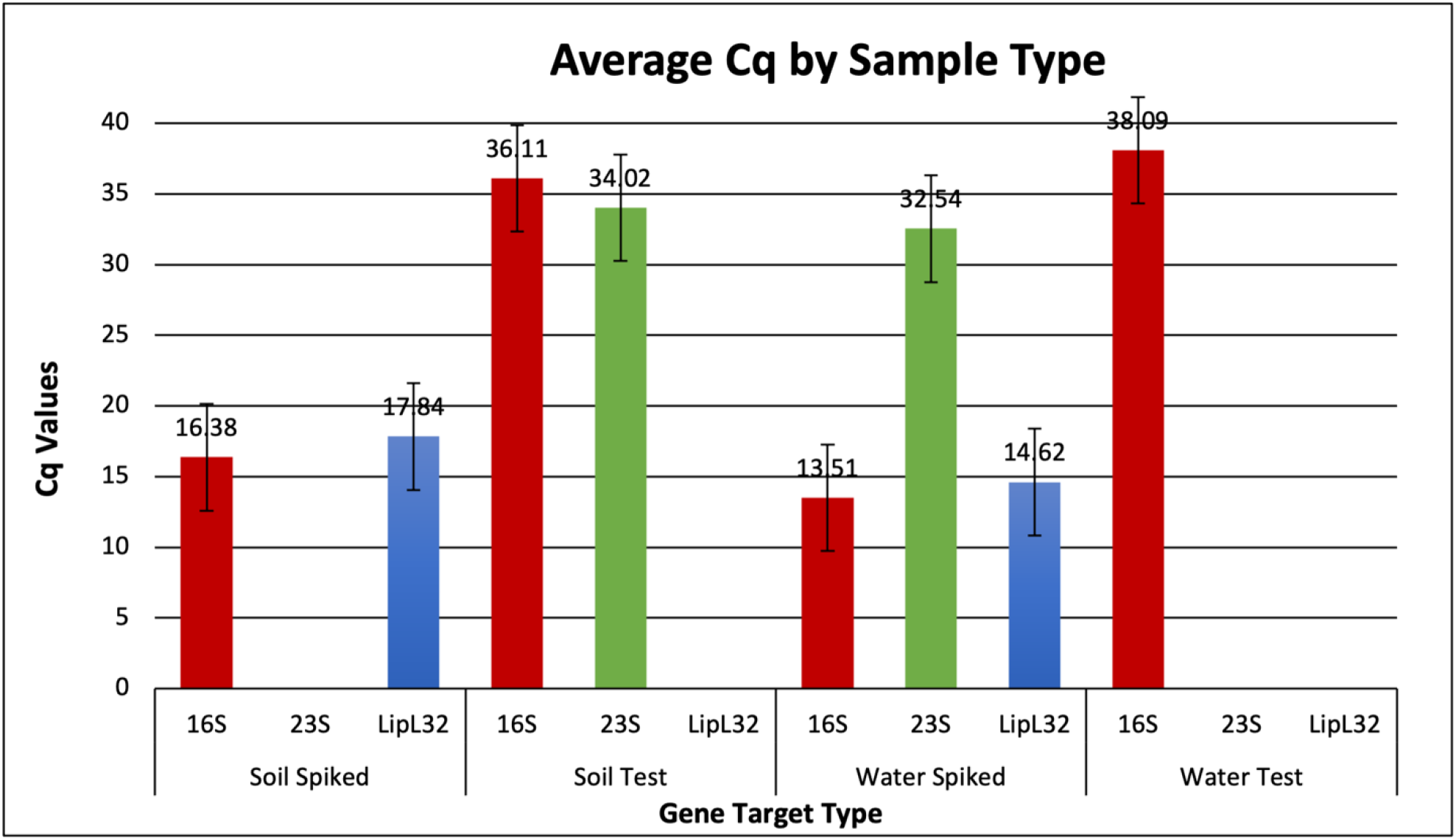
Average Cq values for the water and soil samples that were directly tested with Real-Time PCR. Spiked soil and water samples used as the control for testing and are the only samples where *LipL*32 genes were detected. All samples displayed the presence of *16S rRNA* gene, but only the soil test and water spiked samples had *23SrRNA* genes detected in their sample.

### Culture results

The soil and water samples were cultured in four different types of media with microbial inhibitor combinations. The presence and levels of organisms with morphology compatible with *Leptospira* were recorded with a 0 to +4 ordinal system (Table 1). The cultures with selective antimicrobial inhibitors demonstrated large and earlier increases in bacterial organisms with morphology and motility compatible with *Leptospira* when observed under the DFM. Overall, EMJH cultures with 5-FU or STAFF were favorable for *Leptospira* growth for the water test group. For the soil test samples, the culture results were more variable. The usage of selective antimicrobials in the cultures did not have as much of a visible impact on the growth of *Leptospira* in the soil samples. Overall, Fletcher and EMJH media demonstrated favorable growth for the soil samples, with EMJH performing marginally better than the Fletcher media. All water test and soil test cultures were tested using real-time PCR to confirm the presence of *Leptospira*. For the *23S rRNA* gene marker, all soil and water samples were positive with consistently low Cq values. The cultured test water samples were positive for *LipL32* and *16S rRNA* gene markers. The Cq values for the water samples were consistently around 30 to 35, demonstrating lower levels of *LipL32* and *16S rRNA* in the water samples compared to the cultured soil test samples. The cultured soil test samples had lower Cq values for the *LipL32* and *16S rRNA* gene markers, indicating higher levels of DNA in the soil samples. The growth pattern of *Leptospira-*like organisms in various cultures are shown in supplemental Figure 2

### Sequencing results

The details of results from all sequencing methods are shown in Table 2

**Table 2:** Overall read classification from all the sequencing methods used in this study ^1^-Culture enrichment and metagenomic sequencing; ^2^- Direct metagenomic sequencing; ^3^-16S amplification-based sequencing.

| Source | Total reads | Classified | Chordate | Unclassified | Microbial | Bacterial | Accession |
| --- | --- | --- | --- | --- | --- | --- | --- |
| Water<sup>1</sup> | 233,994 | 69.20% | 0.01% | 30.80% | 69.20% | 69.10% |  |
| Soil<sup>1</sup> | 237,064 | 65.60% | 0.00% | 34.40% | 65.60% | 65.50% |  |
| Water<sup>2</sup> | 7,425 | 83.10% | 0.05% | 16.90% | 83.10% | 82.80% |  |
| Soil<sup>2</sup> | 438,190 | 90.20% | 0.01% | 9.84% | 90.20% | 90% |  |
| Water<sup>3</sup> | 78,756 | 99.38% | 0.00% | 0.62% | 0.00% | 99.38% |  |
| Soil<sup>3</sup> | 68,056 | 99.66% | 0.00% | 0.34% | 0.00% | 99.66% |  |

### Microbial composition profiling and Leptospira classification from 16S dataset

A very low number of reads were classified under the phylum taxon “Spirochaetota” in the water (1 read) and soil (9 reads) samples when 16S rRNA gene sequence dataset was analyzed. The single reads identified in the water sample were closely clustered with the reference *16S rRNA* sequences of two pathogenic *Leptospira spp*., *L. interrogans* and *L. kirschneri*. For the 9 reads obtained from the soil samples, reads were clustered into two separate clusters on the NJ phylogeny. The first cluster was closer to the *16S rRNA* sequences of saprophytic and other environmental *Leptospira* species, while the second cluster was found genetically distant from all *Leptospira* species but closely related to the 16S rRNA of *Leptonema illini* (supplemental figure 3)

**Figure 3.**
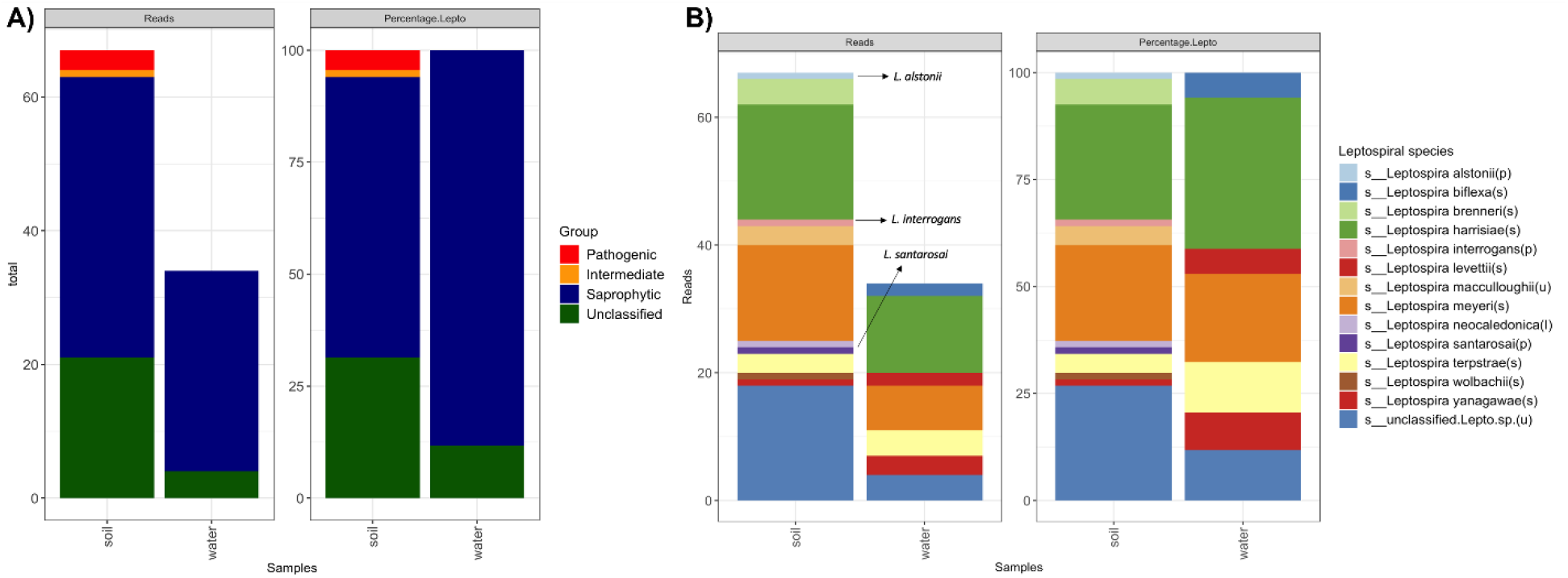
*Leptospira* composition profiles for directly sequenced soil and water samples 3A. Proportion of *Leptospira* clades identified; 3B. *Leptospira* species-level classification. Pathogenic species identified in the soil sample is labeled in the figure. The Group of each *Leptospira* species is annotated in the parenthesis behind species names in the figure legend (p: Pathogenic; i: Intermediate; s: saprophytic; u: Unclassified).

### Microbial composition profiling and Leptospira classification and identification from direct sequencing

A wide range of potentially pathogenic and water-associated microbial sequences were detected from directly sequenced soil (1,438 unique genera) and water (371 unique genera) samples. From those, 102 reads (soil: 67 reads; water: 34 reads) from 12 different *Leptospira sp*. were identified from soil and water samples. Saprophytic *Leptospira spp*. reads were identified in both soil and water samples. Interestingly, pathogenic and intermediate groups of *Leptospira spp*. reads were identified in the soil sample with low coverage. Only three reads of the pathogenic *Leptospira sp*. (1 read from *L. interrogans;* 1 from *L. alstonii*; 1 read from *L*.*santarosai*) and one read of the intermediate *Leptospira sp*. (*L. neocaledonica*) were identified from the soil sample. In addition, around 27% and 12% of *Leptospira* reads identified in the soil and water samples could not be classified at the species level. Figure 3 summarizes microbial classification profiles of direct sequencing results.

### Microbial profiling of the enrichment culture

We pooled positive culture samples from water and soil, prepared a composite sample for each, and proceeded with sequencing. For samples sequenced with culture enrichment, 1,325 unique microbial genera were identified across all samples, with over 60% of all reads classified as bacteria (Table 2). Ninety-eight percent of 30,453 total reads from soil, and 99% of 127,940 total reads from water were classified under *Leptospira*. In total, 34 unique pathogenic and nonpathogenic *Leptospira spp*. were identified in the enrichment cultures. It is interesting to note that 98% and 99% of leptospiral reads in the soil and water samples were classified under either a saprophytic species or an unclassified species. Eleven pathogenic species in the soil from 551 (1.8%) reads and 13 pathogenic species in the water from 859 (0.7%) total reads were identified. In addition, nine intermediate species in soil from 33 reads (0.1%) and ten intermediate species in water from 141 reads (0.1%) were identified. Figure 4 summarizes microbial classification profiles of direct sequencing results.

**Figure 4.**
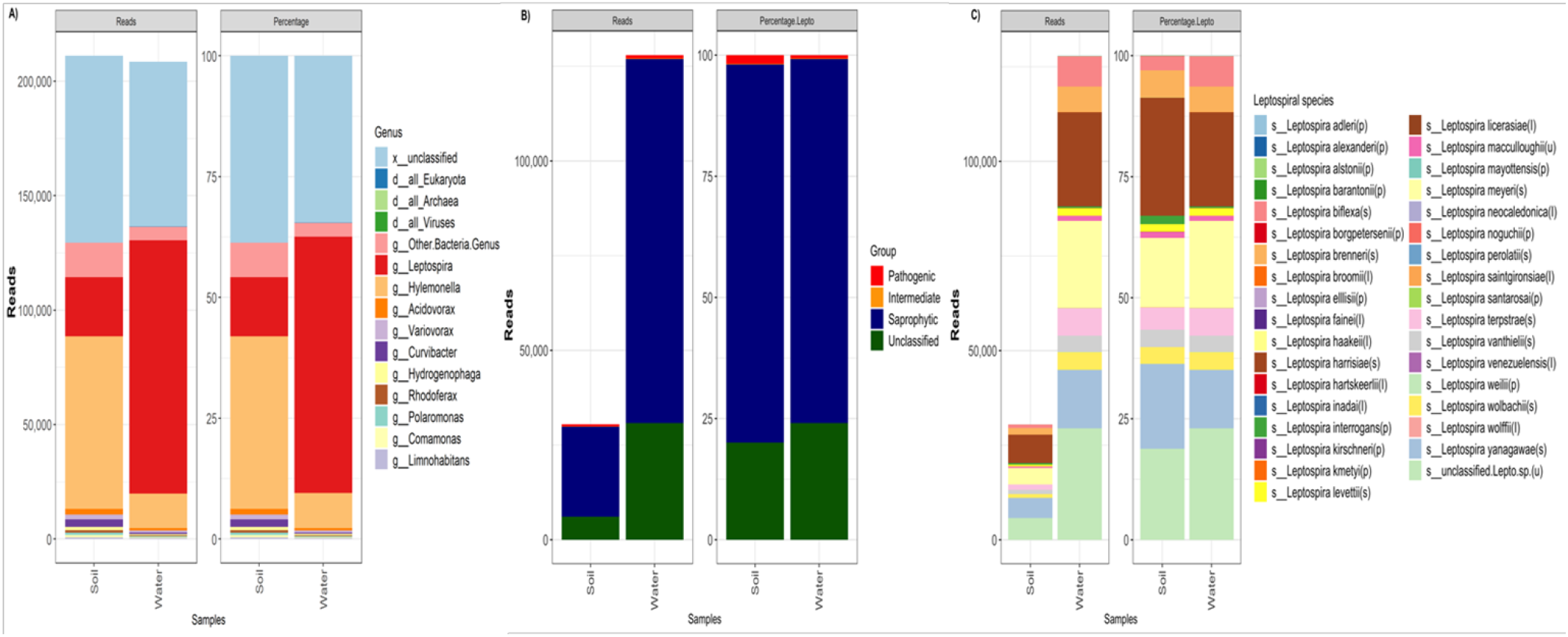
Microbial classification summary statistics of pooled enrichment cultures of soil and water samples. The number and percentage of reads classified under different *Leptospira* species in enrichment cultures sequenced are presented. Group of each *Leptospira* species is annotated in the parenthesis behind species names (p: Pathogenic; i: Intermediate; s: Saprophytic; u: Unclassified). 4A: General microbial profile, 4B: Proportion of *Leptospira* clades identified 4C: *Leptospira* species-level classification.

## Discussion

There is a critical knowledge gap on various aspects of environmental presence, survival, and persistence of *Leptospira*. Exposure to contaminated soil and water is a major risk factor for acquiring leptospirosis in humans and animals. The maintenance of bacteria in the soil and water and its dispersal during extreme weather events may increase the number of cases during such events. Therefore, this work was focused on evaluating and improving *Leptospira* detection from environmental samples. In this study, we observed variations in *Leptospira* detection when different techniques were applied to water and soil samples.

The original *Leptospira* taxonomy divided this genus into two species, pathogenic *L. interrogans* and saprophytic *L. biflexa* based on phenotypic characteristics (1). These two species had numerous serovars based on their serologic reactivity. Later DNA hybridization studies revealed multiple pathogenic species that included group 1 (pathogenic) and group 2 (intermediately pathogenic). Whole-genome sequencing projects further characterized *Leptospira* genomes revealing many genetic attributes that correlate with virulence and pathogenicity (22-24). The presence and the high diversity of *Leptospira* species from soil and water samples from a single location were confirmed in our study. The presence of saprophytic *Leptospira* sequences was confirmed in both water and soil samples in larger proportions. The presence of the pathogenic and intermediate groups was primarily observed in the soil. Amplification of 16S rRNA and sequencing is a very common method used for microbial profiling of environmental samples, however, our data shows that 16S rRNA-based metagenomics may not detect the low-level presence of *Leptospira* in environmental samples. The technique we applied, the enrichment culture followed by metagenomic sequencing, improved the detection of a diverse set of pathogenic and nonpathogenic *Leptospira* in the soil and water samples. Our findings agree with many recent investigations on environmental samples identifying increased diversity of *Leptospira* species in environmental samples (25, 26). These findings emphasize the need to explore the environment as a potential reservoir of pathogenic *Leptospira*. It is important to note that when enrichment cultures followed by sequencing were applied, a diverse population of *Leptospira* could be observed from soil and water samples from a single site. A recent systematic review also supported the presence of *Leptospira* in soil and its dispersion during extreme events of soil disturbance (27). The bacteria may utilize the environmental conditions in the damp soil and may undergo low-level proliferation enabling their persistence in the soil and subsequent transmission to susceptible hosts and hence reservoir animal kidneys are probably not the only source of contamination.

Analyzing environmental samples can be challenging since the sample has increased diversity of organisms present in varying amounts. To study a specific group of organisms in that sample, such as *Leptospira*, methods such as filtration, amplification, and selective culturing can be implemented to remove other environmental organisms that may out-compete and prevent the identification of target bacteria.

A variety of methods are used to detect *Leptospira* from environmental samples. PCR is a widely used method, and multiple gene targets have been evaluated (28). We used three different types of PCR with variable outcomes. PCR directly from soil or water samples did not confirm the presence of pathogenic *Leptospira*. Growing *Leptospira* in culture can be challenging. Our enrichment culture procedure evaluated various media and antimicrobial supplements following sequential filtration and sedimentation for the recovery and detection of *Leptospira*. In previous studies, filtration methods were utilized to accomplish different goals. Some studies used filters that had a large pore size of 0.7 µm to remove or capture bacteria and other studies used filters that were 0.2 µm to capture *Leptospira* on the filter (12, 29). Our methodology used a double filtration system to improve the efficacy. First, we used a 40 µm filter to catch large debris that could block the smaller filter and impede filtration. Then the resulting filtrate was allowed to pass through a 0.45 µm filter to remove larger bacteria, assuming that *Leptospira* with a width of 0.1 µm would pass through the filter. This double filtration method aimed to concentrate *Leptospira* in the samples, increase the chance of recovery, and reduce contamination. Unlike the direct PCR, culture enrichment followed by PCR could detect *Leptospira* DNA in these samples. *Leptospira* culture in the presence of selective antimicrobial inhibitors might have allowed the replication of *Leptospira* while inhibiting major contaminants. Culture enrichment followed by sequencing allowed a better understanding of the diversity of *Leptospira* species present in these samples. Therefore, the sequential application of traditional and molecular methods will improve the pathogen detection and characterization from environmental samples.

Out of the three PCR targets, we used the *16S* primers to amplify DNA from pathogenic and nonpathogenic *Leptospira, Lipl32* primers to amplify DNA from pathogenic *Leptospira*, and 23S primers to amplify DNA from nonpathogenic *Leptospira*. In our study, the direct PCR screening only detected an extremely low amount of saprophytic *Leptospira* DNA from the soil sample and none from the water. Intrinsic differences in amplification efficiencies and the level of original target sequences present in the samples might be a factor that contributed to the lack of detection by our direct PCR methods.

An experimental study on *Leptospira* survival in soil and water microcosms suggested the inability of *Leptospira* to multiply in environmental sites and the environment may be a temporary carrier for the bacteria shed from animal kidneys (30). Interestingly, a recent study experimentally evaluated the suitability of water-logged soil as a medium for *Leptospira* growth (31). They concluded that *Leptospira* can remain in the soil for longer periods in a resting state and proliferate when they come into contact with water. In bodies of water where the soil has not been recently disturbed, pathogenic *Leptospira* may be present but at DNA levels not detectable by direct qPCR. The limit of detection in many studies are based on spiked samples; however, heterogeneity of environmental samples may affect the sensitivity of detection. A detection limit of 10^1^ to 10^2^ leptospires/mL of blood is suggested, but a higher level of *Leptospira* may be required for environmental samples due to a higher level of PCR inhibition, competing bacteria, and DNA degredation from environmental contaminates that might be present in these samples (27, 32, 33). It is worth noting that direct PCR from environmental samples does not validate the presence of viable *Leptospira*. In contrast, enrichment culture followed by PCR or sequencing allows the confirmation of viable bacteria and is potentially a better method for assessing environmental maintenance and transmission risk.

Implementing 16S rRNA amplification allows bacterial DNA in samples to be selectively amplified. This is especially useful in environmental samples, since there can be contamination. Recently, a cost-effective workflow for microbiological profiling using targeted nanopore sequencing of freshwater detected the presence of *Leptospira* (12). We also used a similar method described in this study, and 16S rRNA was amplified using barcoded custom primers followed by Nanopore sequencing. Surprisingly, only a few reads from *Leptospira* spp. were detected by these methods. This could be attributed to the nature of pathogenic *Leptospira* and its propensity to maintain at low levels in the environment or the larger more abundant environmental DNA crowding out pathogenic *Leptospira* strains limiting the number of reads obtained during sequencing. In our study, the direct sequencing from the samples resulted in a greater number of reads compared to 16S rRNA sequencing workflow and allowed better detection of pathogenic *Leptospira*.

Next-generation sequencing allows us to study the abundance and diversity of microbial populations in environmental samples. Long-read sequencing has allowed us to study the complete genomes of organisms that are not culturable or found in association with other organisms in the environment (34). With this technology, the diversity of environmental samples could be captured since the DNA of organisms could be analyzed despite the size or presence of a conserved strain of DNA (35). The DNA of bacteria, protozoa, and animals could all be sequenced from one sample to help investigate the microbiome found in soil, water, and biological fluids. Previously pure, cultured samples had to be used to analyze the genome of an organism, but new metagenomic sequencing technology allows complex contaminated samples to be analyzed. Commercial platforms such as Illumina, Ion Torrent are widely used for this purpose based on short-read sequencing technology, and ONT Nanopore and PacBio sequencing systems use long-read sequencing methods. ONT nanopore method offered us a cost-effective and user-friendly platform without the need for robust equipment. One of limitation of this system is that the pores can become clogged and create a physical barrier. Larger and more prevalent DNA will pass through the pores and possibly clog the pore before the less prevalent genomes can be sequenced and subsequently may lower the sequence output.

Based on our findings, we propose the enrichment culture followed by real time PCR as a point of care test for water surveillance of *Leptospira* presence in the environment and the enrichment culture followed by sequencing to understand the diversity of *Leptospira* species present in these samples. Our future studies will attempt to evaluate optimal sample volume, incubation time, and cost-effectiveness for routine environmental surveillance procedures for the detection and characterization of *Leptospira*. We also anticipate on isolating mixed cultures of *Leptospira* obtained in this study to purity and further characterize the pathogenic species obtained in this study.

## Acknowledgements

We would like to thank the Boheringer and Ingelheim Veterinary Summer Scholar program for providing the opportunity for Myranda Gorman to conduct this research over the summer and UT College of Veterinary Medicine for funding support.

## Supplemental files

**Supplemental figure 1.**
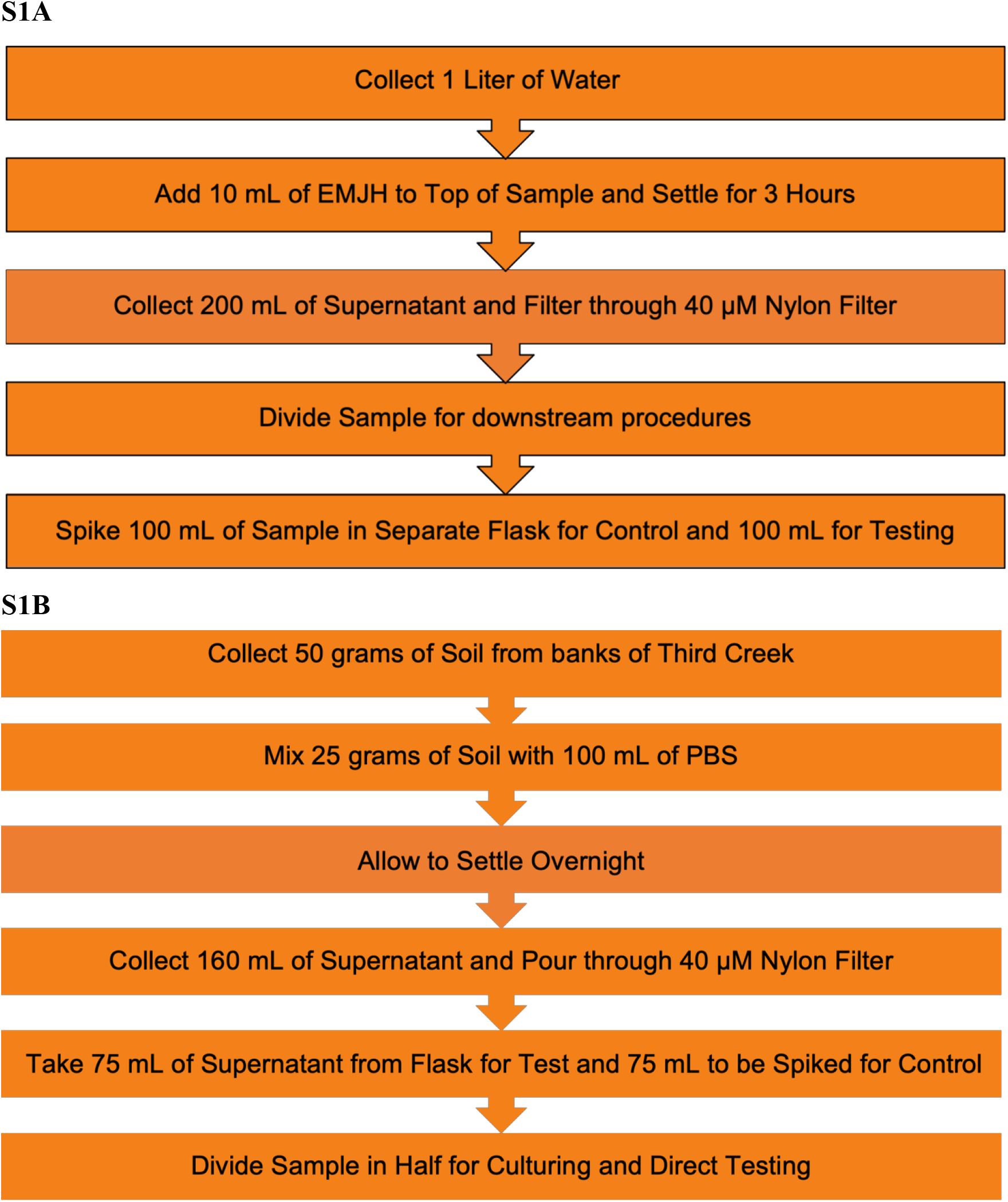
A schematic diagram showing soil and water processing 1A-Water, and 1B-Soil.

**Supplemental figure 2.**
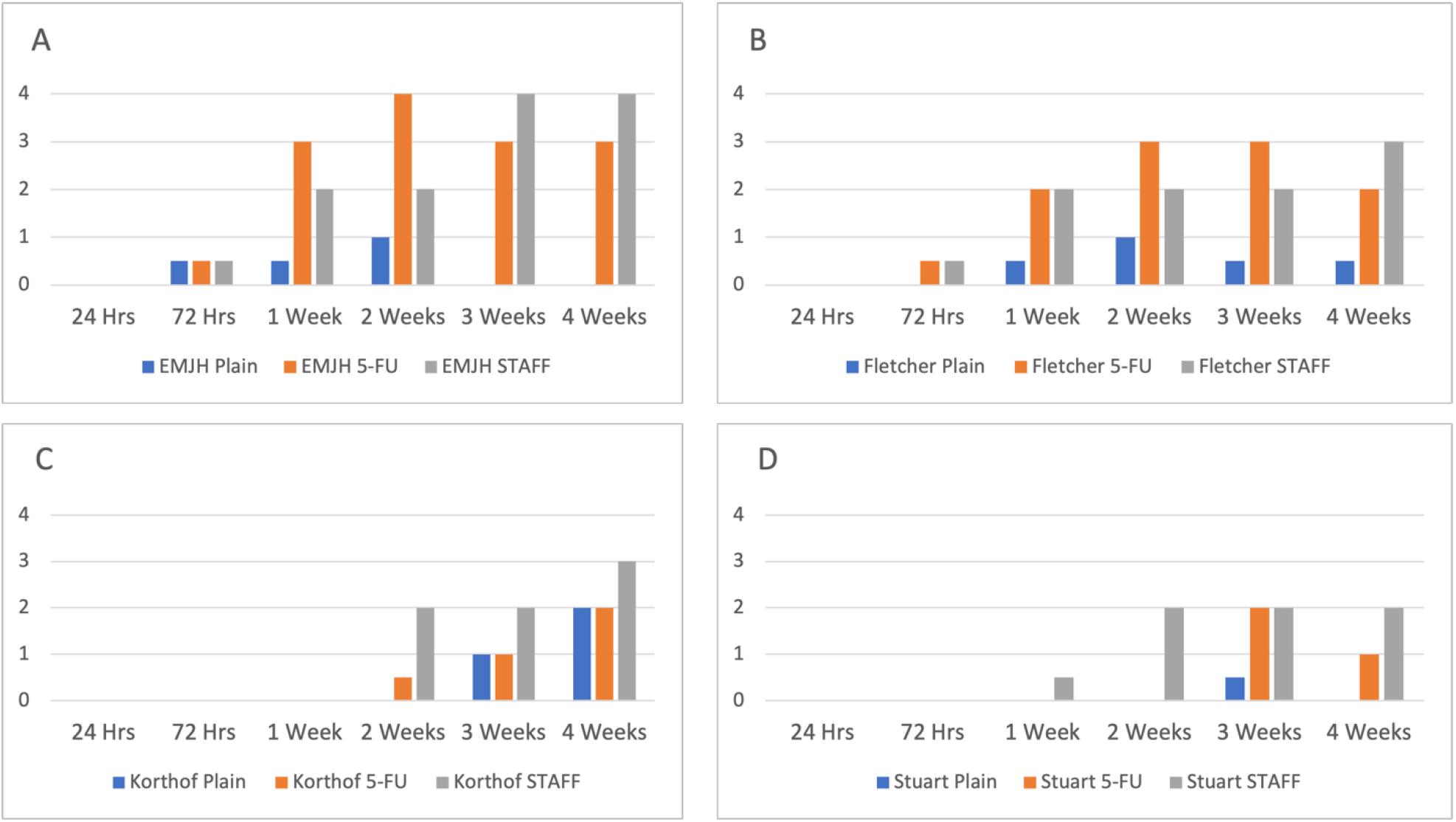
Growth of *Leptospira* like organisms in various cultures. The bar charts grouped here represent the levels (0-4) of *Leptospira* growth in water cultures over a period of four weeks. Each bar chart displays the growth for one of the four medias used along with the different selective antimicrobials added to some cultures. (A: EMJH Media, B: Fletcher, C: Korthof, D: Stuart)

**Supplemental figure 3.**
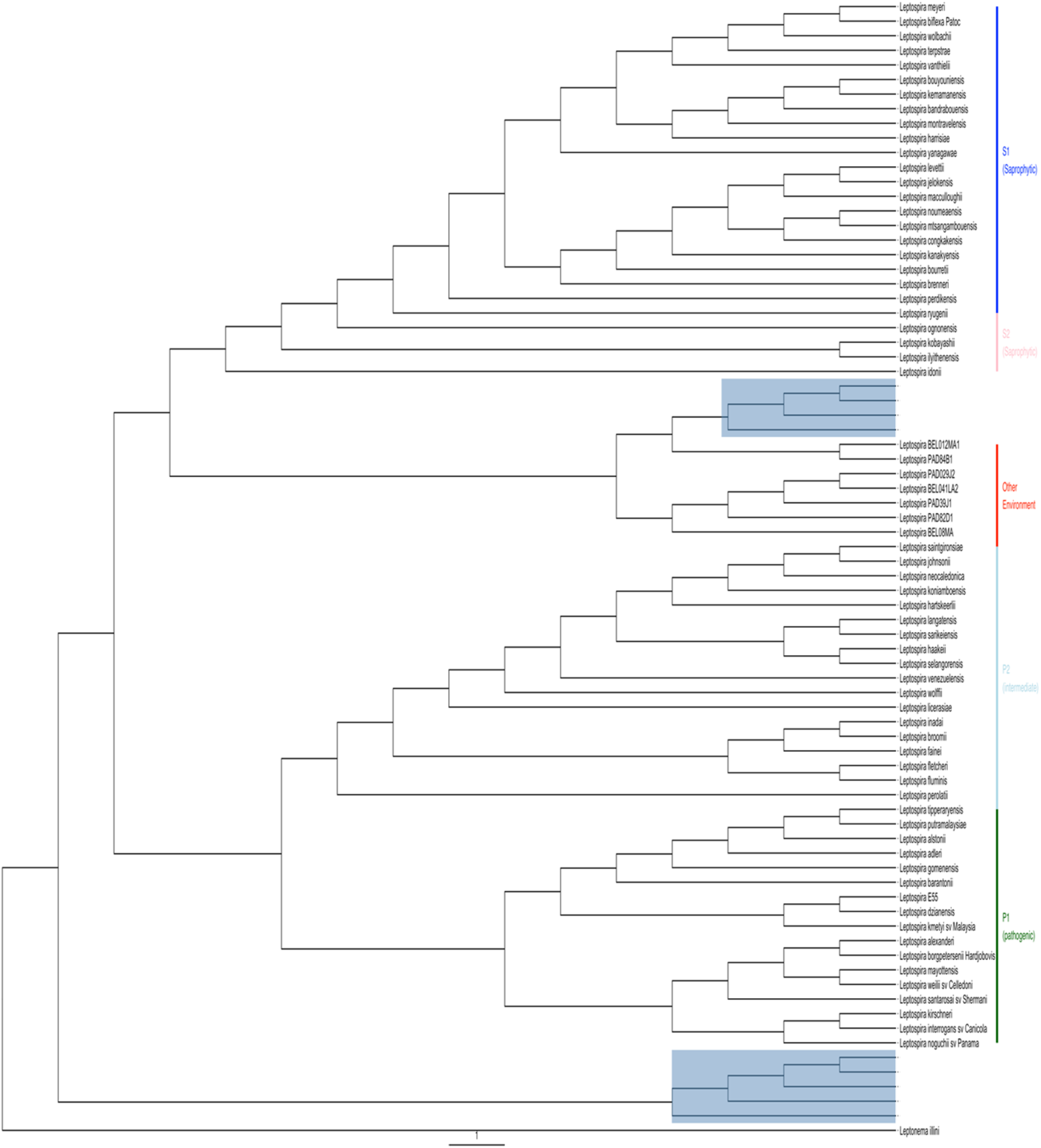
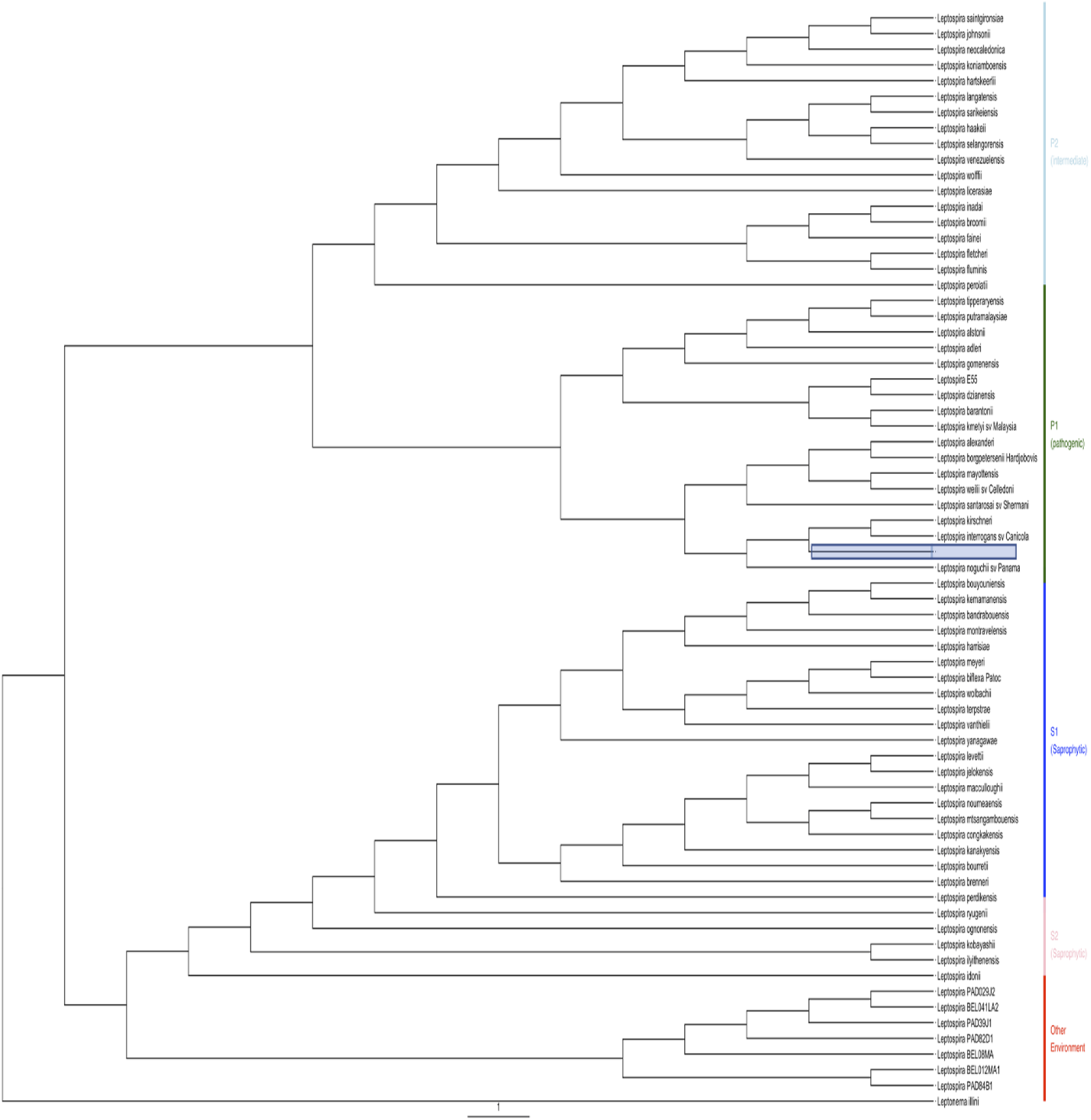
Phylogenetic tree showing position of *Leptospira Classification from 16S dataset*. A-Water samples B-Soil samples

## Notes

### Competing Interest Statement

The authors have declared no competing interest.

